# High throughput protein serial crystallography using a grease matrix and a large-area support film

**DOI:** 10.1101/2025.01.31.635837

**Authors:** Michihiro Sugahara, Saori Maki-Yonekura, Ichiro Inoue, Kiyofumi Takaba, Shun Narai, Hisashi Naitow, Jungmin Kang, Kensuke Tono, Keiji Numata, Tetsuya Ishikawa, Makina Yabashi, Koji Yonekura

## Abstract

Serial femtosecond crystallography (SFX) using ultrashort pulses from X-ray free-electron lasers (XFELs) enables the determination of crystal structures at room temperature while minimizing radiation damage to the samples. This method involves irradiating numerous crystals one by one with XFEL pulses, allowing even the capture snapshots of dynamical structures in biological macromolecules. To achieve this, an efficient sample delivery system is essential for acquiring a large number of diffraction patterns. The most common approach uses a highly viscous grease matrix containing sample crystals, injected into the XFEL path from a narrow nozzle. However, the injection often suffers from clogging issues inside the injector nozzle, resulting in additional challenges such as the need for suitably sized crystals, increased sample consumption and unstable flow rates. Alternatively, a fixed-target approach, which scans a two-dimensional substrate with dispersed samples, can circumvent these issues. However, it must ensure the integrity of biological samples and provide sufficient surface area for efficient data collection. We here present an approach that utilizes a grease matrix and a large-area support film specially designed to address these requirements. This system offers a fast and reliable solution for protein SFX, enabling high-quality structure determination while significantly reducing sample consumption.

## Introduction

Serial femtosecond crystallography (SFX) using X-ray free-electron laser (XFEL) pulses^1–5^ offers a means to overcome typical radiation damage to samples such as proteins^6–14^ and chemical compounds^15–20^. This is achieved through the “diffraction-before-destruction” approach^21^, where numerous crystals are sequentially irradiated with short XFEL pulses at a high repetition rate. Thereby, the SFX has opened new opportunities for ultrafast (femtosecond–microsecond scale) time-resolved studies of structural changes and chemical dynamics^22–36^. Various sample delivery methods for the SFX have been reviewed^37–40^, highlighting the importance of low sample consumption, a stable sample scan rate synchronized with XFEL pulses and prevention of crystal degradation, particularly maintaining the native state of protein samples during the experiments. There are two major types of the sample delivery^37–39^: (i) injection methods^8,41,42^ and (ii) fixed-target methods^12–14,43–45^. The hybrid delivery methods such as tape-drive droplet injector^46^, crystal extractor^47^ and micro-tubing reeling system^48^ have also been introduced.

For protein SFX using the injection method, a liquid jet^41^ containing small crystals is injected at a relatively high speed of ∼10 m sec^-1^, consuming at least 10–100 mg of protein. To reduce the sample consumption, micro-extrusion techniques utilizing viscous carrier media, such as a lipidic cubic phase (LCP)^8,49^, grease^9,50–52^, Vaseline (petroleum jelly)^53^, hydrogels ^54^, *etc*.^55–58^ have been adopted. The micro-extrusion method with these viscous media enables a stable stream at low flow rates of 0.02 –0.6 μl min^-1^, reducing sample consumption to less than ∼1 mg. Thus, viscous carrier media have proven to be highly adaptable for protein SFX, accommodating a wide variety of soluble and membrane proteins^9,11,23,30,34,36,50,51,59–67^. However, the sample injection from a thin injector nozzle often encounters clogging issues with the sample crystals. To avoid clogging during injection, crystal-size filtering is required^37^. Unfortunately, this filtering step approach results in significant sample loss, increasing sample consumption. Moreover, crystal clogging and/or variations in crystal size lead to unstable flow rates (scan speeds) of the sample stream, which often stops diffraction measurement and causes significant difficulties. This is particularly a major obstacle for time-resolved SFX experiments^55^.

The fixed-target method is more straightforward to set up and operate for SFX, as the crystals are simply placed on a substrate such as traditional goniometer-based pins^12,14^, silicon chips^13,43,44,68–70^ or thin films^71–74^, and scanned by XFEL pulses. This method is free of clogging issues, resulting in stable sample scan speeds during the experiment and minimizing measurement dead time, although the scan speed (the data acquisition rate) of fixed-target methods is limited by the motor speeds of the sample stage^13^. For time-resolved crystallography, recent studies have reported structure determination using this method for crystals of proteins^75–78^ and metal organic frameworks (MOFs)^20^. Additionally, it has been demonstrated at a synchrotron from millisecond to second time scale^79^. The fixed-target methods are categorized into two groups^37^: (i) multi-shot goniometer-based approaches^12,14,76^, which expose multiple locations of large single crystals (*e.g*. over ∼50 μm) with controlled rotation, and (ii) multiple small-crystal approaches^13,43,70–72^, which move a sample holder with dispersed small crystals to expose a fresh single crystal to each XFEL pulse. The multiple small-crystal approach is particularly well-suited for SFX experiments, as there are fewer limitations in crystal sizes from a few to more than 50 μm in size. For protein SFX using the multiple small-crystal approach, the use of a sheet-on-sheet (SOS) sandwich structure has been reported to prevent damage caused from crystal dehydration^72^. This structure utilizes nylon mesh^80,81^ and hydrogel media (gelatin and agarose)^73^ to support crystals between the two polyimide films, preventing sinking due to gravity.

Our group has demonstrated a hybrid approach in which a two-dimensional (2D) scan of the substrate coupled with rotation enables highly efficient data collection from small organic molecular crystals^15,17,82,83^. In these studies, we used low-viscosity liquid paraffin to distribute and adhere sample crystals onto a flat-faced polyimide plate with a size of 4 mm × 4 mm. The sample plate was then mounted vertically onto a sample stage with a pin for XFEL exposures. This method is compatible with various crystal types, including small but thick crystals, preferentially oriented plate-like crystals, and crystals ranging from a few to over 50 μm in size, making it particularly well-suited for SFX experiments.

Despite these advances described above, sample preparation for protein SFX remains time-consuming and more efficient, adaptable methods are essential. In this study, we introduce a grease matrix technique into fixed-target protein SFX to further enhance sample delivery. Our method simplifies sample preparation by mixing crystals with a grease matrix medium. Loading the sample involves merely distributing the grease matrix over a large-area PEEK film. The viscous matrix prevents dehydration of the crystals and maintains their distribution by reducing sinking. Crystal orientations within the thick grease layer are random, similar to those in the injection method. Using the grease matrix and a developed sample holder for the PEEK film, we successfully determined the structure of proteinase K from *Engyodontium album* at 1.2 Å resolution.

## Results and discussion

### Sample preparation, data collection and structure analysis

Our previous setup utilized a custom-designed polyimide plate measuring 4 mm × 4 mm in size and 20 to 40 μm in thickness^15,17^, which was scanned in 10 μm steps between sequential XFEL exposures in air, without the use of a vacuum path or helium gas. A larger step would be more suitable for protein crystals that are highly sensitive to radiation^12^, and would require a larger area for efficient data collection and time-resolved studies. This low-background polyimide film is expensive for larger areas, and we tested several materials and found the PEEK film^84^ meets this requirement. Our newly developed sample holder features two windows, each of which provides a 2D scan area of 8.2 mm × 36 mm (**Fig. 1**). Europium (Eu)-derivatised proteinase K crystals measuring 10 μm × 10 μm in size was then mixed with a grease matrix composed of 15% (*w/w*) dextrin palmitate/DATPE grease (DATPE grease)^52^ and uniformly spread over a PEEK film covering the window of the sample holder (**Fig. 2**). The sample holder was scanned in two dimensions at 750 μm sec^-1^, ensuring a 25 μm spacing between XFEL pulses, which operated at a 30-Hz repetition rate at the SACLA facility. This setting allowed for approximately 3 times more points to be shot without human supervision compared to the previous setting [(8,200×36,000/25/25) / (4,000×4,000/10/10) ≈ 2.95]. The wavelength of XFEL was selected as 0.827 Å for efficient SAD phasing, leveraging Eu absorption (*f ’*= -0.08 and *f ’’*=4.76). We collected ∼115,000 diffraction images on a Rayonix MX300-HS detector in about 90 min from the PEEK film over one window (**Fig. 3**). A total sample volume of about 12 μl was used for this experiment, with a crystal number density (after mixing with the matrix) of 1.4 × 10^8^ crystals ml^-1^. We successfully indexed and integrated ∼21,000 images for the proteinase K crystals (space group *P*4_3_2_1_2) yielding a 100% complete dataset with a CC_1/2_ of 0.940 at a resolution of 1.2 Å (**Table 1**).

**Figure 1.**
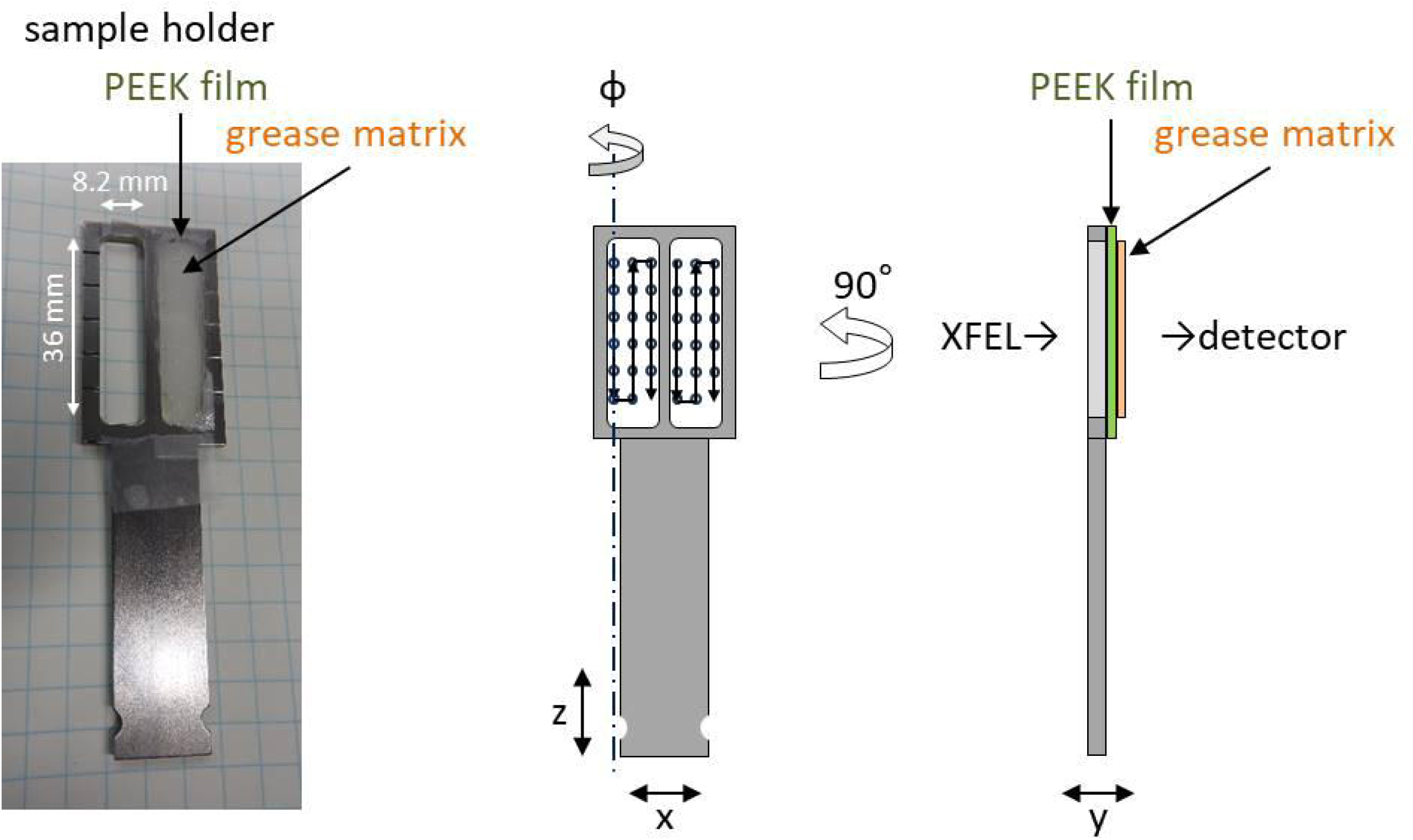
Sample holder. The photograph shows a window with an attached PEEK film coated in a grease matrix. The illustration depicts the 2D scanning of sample windows with XFEL pulses.

**Figure 2.**
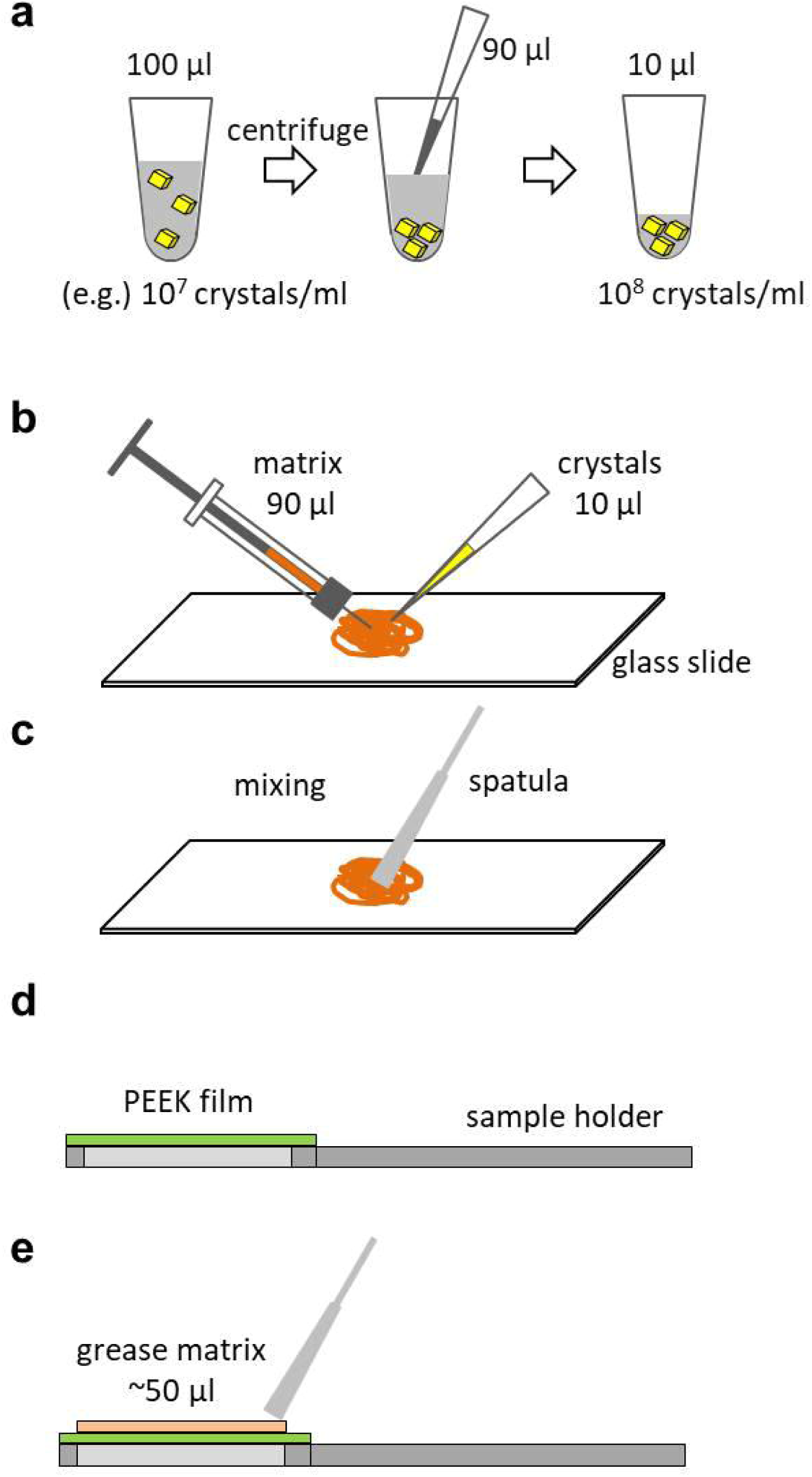
Flowchart of the sample preparation process for the grease matrix technique. (**a**) After centrifuging the crystal sample in the storage solution, the supernatant is removed. (**b, c**) The grease and crystal solution are dispensed onto a glass plate and mixed. (**d, e**) The grease matrix is spread on the PEEK film attached to the sample holder.

**Figure 3.**
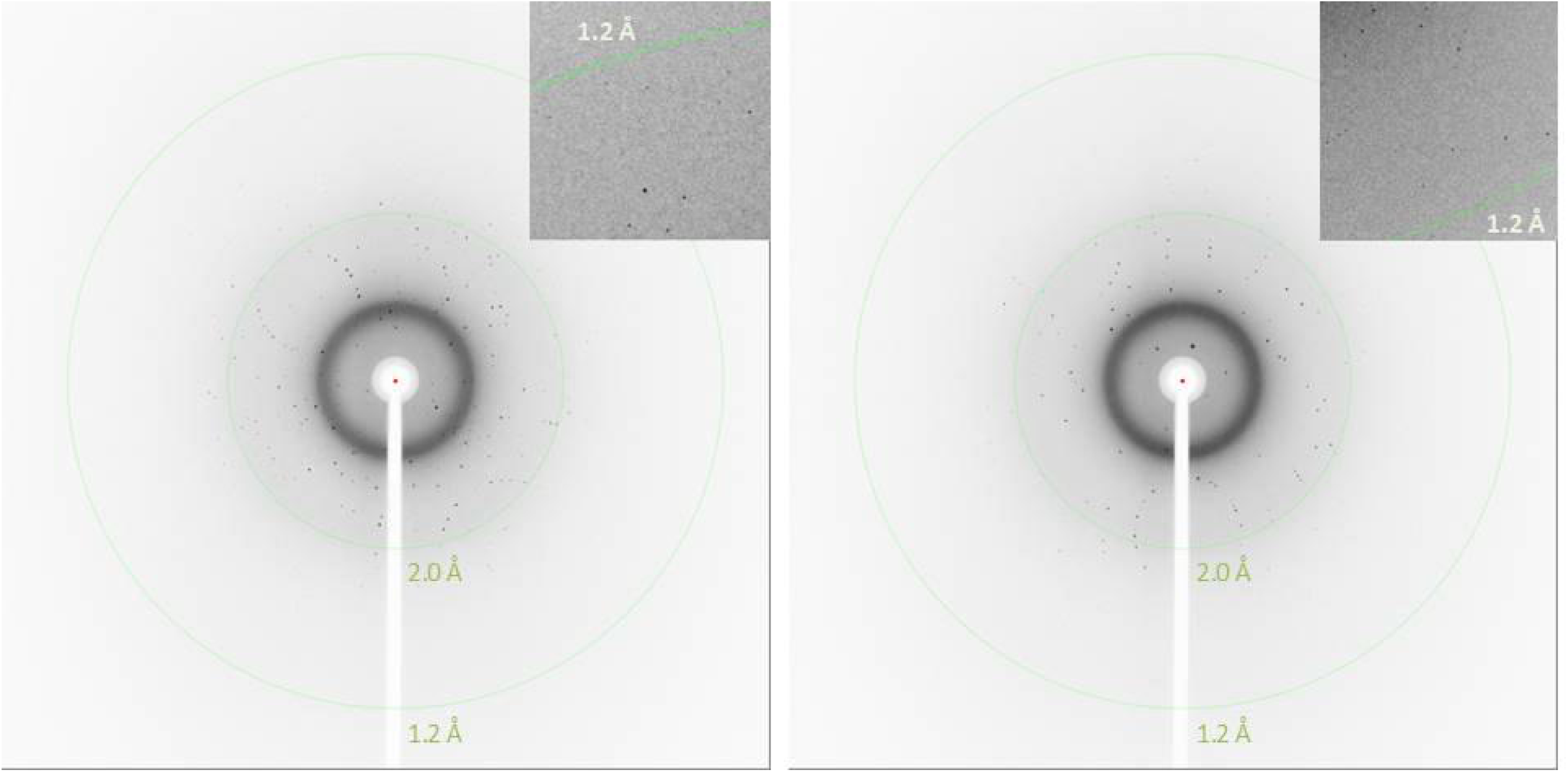
Typical XFEL single diffraction images.

**Table 1.** Crystallographic statistics. Values in parentheses are for the outermost shell.

| **Data collection** |  |
| --- | --- |
| Space group | $P4_32_12$ |
| Unit-cell parameter |  |
| $a = b$ (Å) | 68.70 |
| $c$ (Å) | 108.98 |
| Number of collected images | 114,977 |
| Number of hits | 30,692 (27%) |
| Number of indexed images | 20,763 (68%) |
| Number of total reflections | 21,761,450 |
| Number of unique reflections | 82,104 |
| Resolution range (Å) | 29.6–1.2 (1.22–1.20) |
| Completeness (%) | 100 (100) |
| Multiplicity | 265 (180) |
| $R_{\text{split}}$ (%)<sup>†</sup> | 13.0 (82.0) |
| $CC_{1/2}$ | 0.940 (0.472) |
| $\langle I/\sigma(I) \rangle$ | 6.0 (1.4) |
| **Refinement** |  |
| $R/R_{\text{free}}$ (%) | 16.9/18.3 |
| PDB code | 9kw1 |

In a previous study, the structure of proteinase K was determined using a cellulose hydrogel matrix in the injection method^85^, achieving a resolution of 1.20 Å with ∼82,000 crystals. In this study, we used only ∼21,000 crystals to obtain a dataset at the same resolution of 1.20 Å **(Table 1)**. Data collection statistics indicated no significant discrepancies between the two methods, with a CC_1/2_ of 0.47 and <*I*/*σ*(*I*)> values ranging from 1.3–1.4 in the outermost shell at 1.22–1.20 Å resolution. However, the experimental conditions for this fixed-target study differed from those of the injection case, which employed the cellulose matrix^51^ and the MPCCD detector^86^ at a wavelength of 0.95 Å^85^.

The hit rate (27%) and index rate (68%) achieved in this study were comparable to those of the injection method with crystal carrier matrices^51,52^. Although a ∼50 μl matrix (∼1 mg of protein) was applied to the surface of the PEEK film, we obtained a full dataset by scanning a portion of the film corresponding to a 12 μl aliquot of the matrix (∼0.2 mg protein). In contrast, the continuous sample flow at relative low flow speed (typically 0.02–0.6 μl min^-1^) for the injection method consumes ∼1 mg or less of the sample^42^. Thus, this fixed-target approach can reduce sample amount to 1/3 to 1/5 of that in the injection method. The advantage is even more pronounced when considering crystal-size filtering required in the injection method. The scan speed in both methods appears well-suited for XFEL pulses with repetition rates of 30–60 Hz at the SACLA facility.

### Crystal structure of proteinase K

We then investigated *de novo* phasing from the dataset of Eu-derivatised proteinase K, collected in this study. Single-wavelength anomalous diffraction (SAD) phasing was carried out using *SHELXD* and *SHELXE*^87^. We successfully identified two Eu ions in the asymmetric unit and subsequently solved the substructure (**Fig. 4**). The two Eu-binding sites corresponded to those previously identified for praseodymium (Pr) ions in the proteinase K structure^51,52^. We employed the coordinates of the heavy atoms for both refinement and phase calculation at a resolution of 1.7 Å in *SHEXLE*, which also automatically traced a polyalanine model of proteinase K. Subsequently, *ARP/wARP*^88^ automatically modelled 99% (276 of 279 residues) of the structure with side chains. Finally, we refined the structure at a resolution of 1.20 Å, yielding *R*/*R*_free_ values of 16.9%/18.3%. We were able to observe clear electron density maps of proteinase K (**Fig. 4a**).

**Figure 4.**
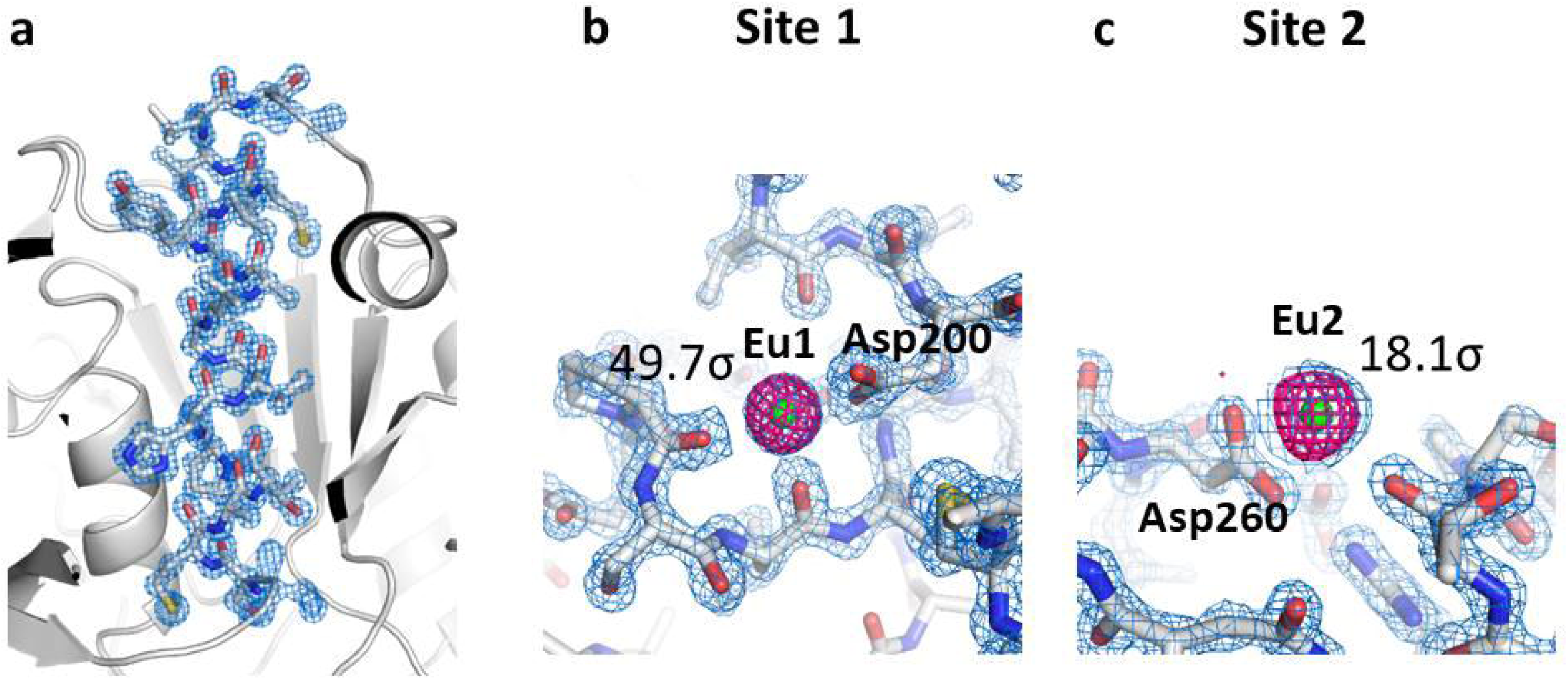
Electron density maps of proteinase K. (**a**) A close-up view of the proteinase K structure with a 2F_o_–F_c_ electron-density map contoured at the 2.0 σ level. This figure was drawn with *PyMol* (http://www.pymol.org). (**b, c**) Close-up views of Eu ion binding sites with 2F_o_–F_c_ electron density maps contoured at the 2.0 σ level (coloured blue). Bound Eu ions are depicted as magenta spheres. The anomalous difference Fourier maps using all indexed images (contoured at the 3.5 σ level) are shown in magenta.The anomalous peak heights obtained from *ANODE* are displayed in figures. These figures were drawn with *PyMol* (http://www.pymol.org).

In the final anomalous difference Fourier maps, generated from all indexed images, we observed significant anomalous peak heights of 49.7 σ and 18.1 σ for the two Eu atoms, calculated using *ANODE*^89^ (**Fig. 4b,c**). Thus, the PEEK film and the grease matrix were found to cause minimal interference with the detection of anomalous signals in protein crystals. In the previous injection SFX study using the DATPE matrix, anomalous scattering signals were detected from the praseodymium (Pr) atoms in the proteinase K structure^52^ from ∼23,000 indexed images. The averaged anomalous densities for the two Pr ions were 46.6 σ for site 1 and 27.9 σ for site 2, according to *ANODE*. Notably, there were no discernible differences in the anomalous difference Fourier maps for the heavy atoms between the two methods. These results indicate that the grease matrix approach utilizing the fixed-target method is effective in detecting weak anomalous signals for de novo phasing in protein SFX data, comparable to the injection method.

### Simplicity and adaptability of the fixed-target procedure with grease matrix

In the fixed-target method, the sample is spread onto a PEEK film, and the sample holder is then mounted on the sample stage for data collection, as described. In contrast, the injection method requires loading the matrix into a sample cartridge, which is then integrated into the sample injector^42^. The injector is positioned on the sample stage and connected to an HPLC pump line that controls the water flow, creating pressure on the plunger within the injector, and to a helium sheath gas line that ensures a stable sample stream. Before data collection, water and gas flows are adjusted to stabilize the stream, which is then aligned with the FEL interaction point. Compared to the injection method, the fixed-target approach involves a significantly simplified sample loading process with fewer procedural steps.

Furthermore, the fixed-target procedure does not need crystal-size filtering. In the injection method, matrix viscosity is critical in maintaining a continuous and stable sample stream from the nozzle of a high-viscosity micro-extrusion injector during serial sample loading. A viscosity-adjustable matrix is necessary to maintain a stable sample stream for the injection method^52^. In contract, the fixed-target method does not demand precise adjustments to matrix viscosity. However, a low-viscosity matrix may cause crystal sinking on the PEEK film^73^. In this case, we employed a 15% DATPE grease matrix to keep the crystals in fixed positions during data collection.

Our fixed-target sample holder is compatible with the capacity for sample rotation, based on a previously developed model^15,17^. In this study, however, we collected a data set without rotation. Proteinase K crystals sized 10 μm were randomly distributed within a matrix of ∼200 μm thickness. Crystals smaller than 30 μm are expected to exhibit random orientation within a matrix of ∼100 μm thickness^9^. For crystals with preferential orientations, the sample stage is initially aligned to the eucentric position and subsequently tilted from 60° to 0° around the phi axis during XFEL exposure (**Fig. 1**)^17^. Thus, this approach offers high adaptability and has the potential to improve the acquisition of high-quality data from various challenging targets.

## Conclusion

This study has introduced a fixed target protein SFX method, utilizing the grease matrix and the large-area support film. This approach significantly reduces the time and labour required for sample preparation and loading, while minimizes sample consumption. It is also demonstrated that neither the matrix nor sample support film interfere with detection of anomalous signals in protein crystals, enabling high-quality structure analysis. This adaptation enhances the versatility of the technique, making it applicable to a broad range of samples, including not only organic small molecular crystals but also protein crystals. This method will be well-suited for time-resolved studies using XFEL and synchrotron X-ray sources.

## Materials and methods

### Sample preparation

We prepared proteinase K (No. P2308, Sigma) crystals sized 10 μm × 10 μm following the previously reported protocols^50,90^. After centrifuging a 100-μl crystal sample of the storage solution (with a crystal density of 4.8 × 10^7^ crystals ml^−1^) at ∼1,300 *g*–3,000 *g* for 10 s using a compact tabletop centrifuge, we removed a 90-μl aliquot of the supernatant. Subsequently, we added a 10-μl sample of the concentrated crystal solution to a 90-μl heavy-atom solution, comprising 27.8 mM EuCl_3_ (HR2-450-8, Hampton Research), 0.5 M NaNO_3_, and 0.1 M MES–NaOH (pH 6.5). We then incubated the crystal solution at 20 °C for 90 min. After centrifuging a 100-μl crystal sample of the heavy-atom solution for 10 s, we removed a 90-μl aliquot of supernatant (**Fig. 2a**). For the grease matrix, we employed a 15% (*w/w*) dextrin palmitate/dialkyl tetraphenyl ether oil (DATPE) grease^52^. We dispensed a 10-μl aliquot of the crystal suspension into 90 μl of 15% (*w/w*) dextrin palmitate/DATPE grease on a glass slide, then mixed them with a spatula (**Fig. 2b,c**). A polyether ether ketone (PEEK) film (Urban Crown Co., Ltd., 2000-012G, thickness 12 μm)^84^ was affixed to the sample holder window (**Fig. 2d**). A ∼50 μl aliquot of the matrix was evenly spread over the film using a spatula (**Fig. 2e**).

### Data collection

We carried out the experiments at SACLA^3,91^ using femtosecond X-ray pulses at a wavelength (photon energy) of 0.827 Å (15 keV) with a pulse energy of ∼200 μJ. Each X-ray pulse delivered ∼10^12^ photons within a 10-fs duration (FWHM) to the sample. The X-ray beam was focused via Kirkpatrick-Baez mirrors^92^ to achieve a spot size of 1.5 μm × 1.5 μm. We collected diffraction data on BL2 in the experimental hatch 3 of the SACLA XFEL facility^93^. The crystals were kept at approximately 25 °C in the experimental hatch. Diffraction images were recorded at 30 Hz through 2D scanning of the matrix sample using a Rayonix MX300-HS detector operating in 4-by-4 binning mode (**Fig. 1**). The sample stage allowed for movement along the XYZ axes and rotation around the phi axis^15,17^. The sample plate was scanned with XFEL pulses by moving the stage in the XZ plane at a speed of 750 μm s^-1^ (25 μm × 30 Hz).

### Structure determination

Bragg spot-containing image files were identified using the diffraction data processing program *DIALS* version 3.5.0^94^. Frames exhibiting more than 10 identified spots were converted to the HDF format using *Python* with the h5py package, and subsequently processed with the *CrystFEL* suite version 0.10.2 for indexing and integration of the intensities^95^. We followed procedures adapted for the SACLA data acquisition system^96,97^. We determined diffraction peak positions using the *peakfinder8* algorithm^98^ and passed them to *MOSFLM*^99^ or *DirAx*^100^ for indexing. We applied no sigma cutoff or saturation cutoff. We merged the measured diffraction intensities using the *process_hkl* in the *CrystFEL* suite^95^ with scaling (--*scale* option). For the Eu-derivatised proteinase K, we carried out substructure search, phasing, and phase improvement using the *SHELX C, D,* and *E* programmes^87^. We fed the auto-traced model from *SHELXE* into *ARP/wARP*^88^ in the *CCP4* suite^101^. We performed manual model revision and structure refinement using *Coot*^102^ and *PHENIX*^103^, respectively. In **Table 1**, we summarise details of the data collection and refinement statistics.

## Acknowledgements

The XFEL experiments were carried out at the BL2 of SACLA with the approval of the Japan Synchrotron Radiation Research Institute (proposal no. 2023B8020). The authors thank the SACLA beamline staff for technical assistance. This research was partly supported by RIKEN Engineering Network Program, the Platform Project for Supporting Drug Discovery and Life Science Research (Basis for Supporting Innovative Drug Discovery and Life Science Research) from AMED under Grant Number JP24ama121006, and JST-Mirai Program Grant Number JPMJMI23G2.

